# Genes influenced by MEF2C contribute to neurodevelopmental disease via gene expression changes that affect multiple types of cortical excitatory neurons

**DOI:** 10.1101/2019.12.16.877837

**Authors:** Donna Cosgrove, Laura Whitton, Laura Fahey, Pilib Ó Broin, Gary Donohoe, Derek W. Morris

**Author notes:** These authors contributed equally to this work (Donna Cosgrove, Laura Whitton). Corresponding author: Dr Derek Morris, Room 106, Discipline of Biochemistry, National University of Ireland Galway, University Road, Galway, Ireland.

## Abstract

Myocyte enhancer factor 2 C (MEF2C) is an important transcription factor during neurodevelopment. Mutation or deletion of *MEF2C* causes intellectual disability (ID) and common variants within *MEF2C* are associated with cognitive function and schizophrenia risk. We investigated if genes influenced by MEF2C during neurodevelopment are enriched for genes associated with neurodevelopmental phenotypes, and if this can be leveraged to identify biological mechanisms and individual brain cell types affected. We used a set of 1,052 genes that were differentially expressed in the adult mouse brain following early embryonic deletion of *Mef2c* in excitatory cortical neurons. Using GWAS data, we found these differentially expressed genes (DEGs) to be enriched for genes associated with schizophrenia, intelligence and educational attainment but not autism spectrum disorder (ASD). Using sequencing data from trios studies, we found these DEGs to be enriched for genes containing *de novo* mutations reported in ASD and ID, but not schizophrenia. Using single cell RNA-seq data, we identified that a number of different excitatory glutamatergic neurons in the cortex were enriched for these DEGs including deep layer pyramidal cells and cells in the retrosplenial cortex, entorhinal cortex and subiculum. These data indicate that genes influenced by MEF2C during neurodevelopment contribute to cognitive function and risk of neurodevelopmental disorders. Within excitatory neurons, common SNPs in these genes contribute to cognition and SZ risk via an effect on synaptic function based on gene ontology analysis. In contrast, rare mutations contribute to earlier onset ASD and ID via an effect on cell morphogenesis.

**Author Summary:** Schizophrenia is a complex disorder caused by many genes. Current drugs for schizophrenia are only partially effective and do not treat cognitive deficits, which are key factors for explaining disability. Here we take an individual gene identified in genetic studies of schizophrenia and cognition called *MEF2C*, which on its own is a very important regulator of brain development. We use data from a mouse study where MEF2C has been stopped from functioning or knocked out during brain development. The effect of that knock out has been measured when the mice reach adulthood, in the form of a set of differentially expressed genes (DEGs) from the somatosensory cortex. We found that this set of DEGs contains more genes than expected by chance that are associated with schizophrenia and cognition or contain rare new (*de novo*) mutations reported in autism and intellectual disability. Using gene expression data from single brain cells, we identified that a number of specific excitatory glutamatergic neurons in the cortex were enriched for these DEGs. This study provides evidence that the molecular mechanisms that underpin schizophrenia and cognitive function include disruption of cell types influenced by MEF2C.

## Introduction

Myocyte enhancer factor 2 (MEF2) is a transcription factor with a highly conserved DNA binding domain and is crucial for differentiation and development through potentiation of other regulatory mechanisms. MEF2C is required for neurogenesis (1), neuronal distribution, and electrical activity in the neocortex (2, 3). MEF2C is the earliest of four subclasses of Mef2 transcription factors to be expressed during embryonic brain development and maintains high levels of expression in adult neurons (4). The N-termini of MEF2 contain a highly conserved MADS-box, with an adjacent ‘MEF2 domain’ that mediates DNA binding and other interactions; and the C-terminal regions display patterns of alternative splicing. The activity of MEF2 proteins is also tightly regulated by class II histone deacetylases (HDACs) interacting directly with the MADS domain (5). This interaction with HDACs indicates a point of convergence of multiple epigenetic regulatory mechanisms, (2) with MEF2C playing a role in epigenetic alterations in chromatin configuration. This includes transmitting extracellular signals to the genome and activating genetic programmes that control cell differentiation, proliferation, survival and apoptosis.

Although *Mef2c* homozygous knockout mice do not survive the gestation period due to incomplete cardiac morphogenesis (2, 6), *Mef2c* heterozygous mice are viable, although only 44% survive until three weeks, compared to wild-type mice. *Mef2c* conditional knockout mice exhibit reduced spatial learning and memory function (1) and deficits in locomotor activity and motor coordination (7). *Mef2c* is a negative regulator of both embryonic and postnatal synaptogenesis (7, 8), with heterozygous knockout mice displaying neuronal and synaptic abnormalities (9). This includes a significant increase in dendritic spine density in the hippocampus, seen in conditional knockout mice that targeted *Mef2c* deletion selectively to the brain during embryogenesis and postnatally (7), and during CNS deletion of *Mef2c* (10). Adult hippocampal slices from Mef2c-null brains compared with WT showed characteristics of an immature neuronal network (1), and abnormal neuronal behaviour was observed in the neocortex during development. This indicates that MEF2C plays a pivotal role in early neuronal differentiation (11). MEF2C specific knockout impairs hippocampal-dependent learning and memory by increasing synapse number and potentiating synaptic transmission. *Mef2c* expression is normally upregulated during neuronal differentiation and maturation (10, 12). However, deletion of *Mef2c* in the brain postnatally did not impact learning and memory, measurements of synaptic plasticity, or several behavioural measures suggested to recapitulate aspects of autism spectrum disorders in mice (7), implying a distinct role for MEF2C in the prenatal versus the postnatal brain.

Harrington et al. (13) sought to evaluate the role of MEF2C in neurodevelopment by generating conditional knockout (cKO) of *Mef2c* in forebrain excitatory neurons of mice, and investigating the downstream effects on cortical synaptic transmission. The effect observed was a small reduction in glutamatergic synaptic transmission and a large increase in inhibitory synaptic transmission, with MEF2C directly regulating the densities of both excitatory and inhibitory synapses. Data indicated that endogenous Mef2c functions predominantly as a transcriptional repressor to inhibit target genes that promote excitatory synapse elimination and inhibitory synapse formation and/or stability. Differentially expressed genes (DEGs) from the somatosensory cortex of *Mef2c* cKO vs control mice were enriched for genes involved in neuron differentiation and development (up-regulated genes) and synaptic transmission and ion transport (down-regulated genes). This set of DEGs was enriched for autism spectrum disorder (ASD) risk genes. Finally, analysis of *Mef2c* cKO mice identified impairments in multiple behavioural phenotypes, e.g. fear learning and memory, multiple social behaviours, socially-motivated ultrasonic vocalizations, reward-related behaviours and repetitive motor behaviours. These overlap with human neurodevelopmental disorders such as ASD, intellectual disability (ID) and schizophrenia (SZ) where cognitive function is impaired (13).

A *MEF2C* haploinsufficiency syndrome has been identified where affected individuals are frequently non-verbal (14), with severe intellectual disability, epilepsy, stereotypic abnormal movements, minor dysmorphisms, and brain abnormalities (15–17). The relevance of *MEF2C* to SZ and cognition is highlighted by a study showing that (a) MEF2C binding motifs are enriched within the pool of top genome-wide association studies (GWAS) SNPs for schizophrenia, (b) there is a specific over-representation of MEF2C motifs among sequences with SZ-associated histone hypermethylation based on chromatin profiling in neuronal nucleosomes extracted from prefrontal cortex of cases and controls, and (c) *Mef2c* upregulation in mouse prefrontal projection neurons consistently resulted in enhanced cognitive performance in working memory and object recognition paradigms (18).

A number of very large GWAS have recently been published for ASD (19), SZ (20), cognitive ability / human intelligence (IQ) (21–23) and educational attainment (EA) (24, 25). These studies identify very strong association between *MEF2C* and IQ and EA in the general population, plus association for *MEF2C* with SZ but not with ASD. Based on *MEF2C* association with cognition and SZ risk, our hypothesis was that genes differentially expressed upon cKO of *Mef2c* during neurodevelopment would also be associated with these phenotypes. We investigated this using the set of DEGs reported by Harrington et al. (13). We tested and found that the DEGs are enriched for genes associated with IQ, EA and SZ using gene-set analysis of GWAS data and are enriched for genes containing *de novo* mutations reported in exome sequencing studies of ASD and ID trios. To gain further biological insights, we then leveraged these contributions of the MEF2C-influenced genes to neurodevelopment to investigate which individual cell types are enriched for these genes using single cell RNA-seq (scRNA-seq) data from the mouse brain and performed gene ontology analysis to help identify the molecular mechanisms involved.

## Results

### Comparison of association signals at MEF2C in GWAS data

We reviewed the largest published GWAS, which report that SNPs at the *MEF2C* locus are genome-wide significant for SZ, IQ and EA but not ASD (S1 Figure A-D). The top associated SNPs for IQ (rs34316) and EA (rs254781) are in high linkage disequilibrium (LD; *r*^*2*^=0.98) and span the 3’ end and downstream region of the gene. Another SNP, rs639725, which is in high LD with these IQ and EA associated SNPs (*r*^*2*^>0.98), has been reported as an expression quantitative trait locus (eQTL) for MEF2C in the cerebellum of Alzheimer’s disease patients and healthy controls (26) (identified via HaploReg v4.1), where expression quantitative trait locus (eQTL) analysis indicates that the alleles associated with reduced IQ and EA are associated with increased expression of *MEF2C*. We did not find eQTLs related to these GWAS SNPs in the other datasets investigated that included both adult and fetal samples. The top associated SNP for SZ is also in the downstream region but not in high LD with the IQ and EA associated SNPs indicating that there are different genetic variants influencing SZ risk and cognition. We found no evidence of an eQTL involving the SZ risk SNP. *Post mortem* gene expression analysis of cerebral cortex reports an increase in MEF2C expression in SZ cases compared to controls (27). The effect of the SZ risk SNP at MEF2C remains to be elucidated but these *post-mortem* gene expression results indicating that increased expression of MEF2C is observed in SZ is consistent with the eQTL data that points to increased expression of MEF2C being associated with reduced IQ and EA.

### Analysis of MEF2C cKO gene-set in GWAS data

We used a set of 1,055 DEGs (listed in S1 Table) based on an RNA-seq study that captured the transcriptional changes in adult male mouse brain that result from the early embryonic deletion of *Mef2c* in cortical and hippocampal excitatory neurons (13). RNA-seq was performed on mRNA isolated from the somatosensory cortex of *Mef2c* cKO or control littermates (13). These genes that were differentially expressed upon cKO of MEF2C were enriched for genes associated with SZ (P=1.70×10^−06^), IQ (P =9.88×10^−08^) and EA (P=2.24×10^−10^) but not ASD (P=0.247; see Table 1). Brain-expressed genes are a major contributor to these phenotypes. It is possible that the enrichment detected here could be due to the MEF2C gene-set representing a set of brain-expressed genes. However, the MEF2C enrichment was robust to the inclusion in the analyses of both ‘brain-expressed’ (n=14,243) and ‘brain-elevated’ (n=1,424) gene-sets as covariates (S2 Table). To examine if the enrichment we detect for SZ, IQ and EA is a property of polygenic phenotypes in general, we obtained GWAS summary statistics for ten phenotypes and tested the MEF2C gene-set for enrichment in each one. These were a child-onset psychiatric disorders, other brain-related disorders, non-brain related diseases, and height. No enrichment was detected for any of the ten phenotypes (S3 Table).

**Table 1:** MAGMA gene-set analysis of the MEF2C gene-set

| Phenotype | Gene-set | Gene N | Beta | SE<sup>a</sup> | P value<sup>b</sup> |
| --- | --- | --- | --- | --- | --- |
| ASD | All DEGs | 1028 | 0.019 | 0.027 | 0.247 |
| EA | All DEGs | 1028 | 0.205 | 0.043 | **$1.17 \times 10^{-06}$** |
| IQ | All DEGs | 1050 | 0.202 | 0.037 | **$9.88 \times 10^{-08}$** |
| SZ | All DEGs | 1033 | 0.131 | 0.035 | **$8.21 \times 10^{-05}$** |
| ASD | Down reg. genes | 578 | 0.024 | 0.036 | 0.249 |
| EA | Down reg. genes | 578 | 0.227 | 0.056 | **$2.32 \times 10^{-05}$** |
| IQ | Down reg. genes | 587 | 0.216 | 0.051 | **$1.23 \times 10^{-05}$** |
| SZ | Down reg. genes | 577 | 0.162 | 0.046 | **0.00019** |
| ASD | Up reg. genes | 450 | 0.009 | 0.039 | 0.407 |
| EA | Up reg. genes | 450 | 0.148 | 0.063 | 0.010 |
| IQ | Up reg. genes | 463 | 0.162 | 0.055 | **0.0019** |
| SZ | Up reg. genes | 456 | 0.078 | 0.050 | 0.062 |

<sup>a</sup> SE = Standard Error<sup>b</sup> P-values in bold survived Bonferroni multiple test correction for the 12 tests.

Gene ontology enrichment analysis of this gene-set had identified that the up-regulated genes were likely involved in neuron differentiation and development whereas the down-regulated genes were involved in synaptic transmission and ion transport (13). To investigate if the up- and down-regulated genes made different contributions to SZ, IQ and EA, we performed gene-set analysis on the up-regulated and down-regulated gene sets separately. Overall, the down-regulated gene-set showed stronger enrichment for SZ, IQ and EA than the up-regulated gene-set (see Table 1).

### Analysis of MEF2C gene-set using data on *de novo* mutations

To investigate the contribution of rare variants in the MEF2C gene-set to SZ, ASD and ID, we tested if the gene-set was enriched for *de novo* mutations (DNMs) that have been reported in trios-based studies of these disorders. We detected strong enrichment for LoF DNMs reported in ASD patients in the MEF2C gene-set where we observed 45 DNMs in the set of DEGs when expecting just 24 DNMs (enrichment value of 1.88; P=7.96×10^−05^). We also found enrichment for missense DNMs in the ASD trios but this did not survive multiple test correction. We detected strong enrichment for missense DNMs reported in ID patients in the MEF2C gene-set (P=3.57×10^−06^; see Table 2) where the number of 25 observed DNMs was nearly 3-fold higher than expected. The sample size for ID (n=192 trios) was small for observing a significant enrichment of LoF DNMs. As the common variant enrichment signal was stronger in the down-compared to the up-regulated genes, for follow-up we tested for enrichment of LoF DNMs in ASD patients and missense DNMs in ID patients in these subsets of genes separately. For both disorders, we found the DNMs tested to be similarly significantly enriched in both down- and up-regulated genes (data not shown). The genes containing LoF DNMs in ASD cases and missense DNMs in ID cases are identified in S1 Table. By way of control, the MEF2C gene-set was not enriched for synonymous DNMs (unlikely to be functional) reported in ASD or ID trios. In addition, both control trios and trios including unaffected siblings of ASD patients showed no enrichment for DNMs in the MEF2C gene-set (Table 2).

**Table 2:** De novo mutation analysis of the MEF2C gene-set

| **Autism Spectrum Disorder (3,985 trios)** |  |  |  |  |
| --- | --- | --- | --- | --- |
| **Mutation class** | **# Observed Mutations** | **# Expected Mutations** | **Enrichment value** | **P value<sup>a</sup>** |
| Silent | 69 | 81.2 | 0.85 | 0.924 |
| Missense | 212 | 177.2 | 1.2 | 0.0060 |
| LoF | 45 | 24.0 | 1.88 | **7.96x10<sup>-05</sup>** |
| **Intellectual Disability (192 trios)** |  |  |  |  |
| **Mutation class** | **# Observed Mutations** | **# Expected Mutations** | **Enrichment value** | **P value<sup>a</sup>** |
| Silent | 4 | 3.9 | 1.02 | 0.549 |
| Missense | 25 | 8.5 | 2.93 | **3.57x10<sup>-06</sup>** |
| LoF | 2 | 1.2 | 1.73 | 0.321 |
| **Schizophrenia (1,024 trios)** |  |  |  |  |
| **Mutation class** | **# Observed Mutations** | **# Expected Mutations** | **Enrichment value** | **P value<sup>a</sup>** |
| Silent | 19 | 20.9 | 0.911 | 0.688 |
| Missense | 49 | 45.5 | 1.08 | 0.322 |
| LoF | 5 | 6.2 | 0.812 | 0.735 |
| **Unaffected siblings (1,995 trios)** |  |  |  |  |
| **Mutation class** | **# Observed Mutations** | **# Expected Mutations** | **Enrichment value** | **P value<sup>a</sup>** |
| Silent | 39 | 40.7 | 0.959 | 0.623 |
| Missense | 93 | 88.7 | 1.05 | 0.338 |
| LoF | 18 | 12.0 | 1.5 | 0.0627 |
| **Controls (54 trios)** |  |  |  |  |
| **Mutation class** | **# Observed** | **# Expected** | **Enrichment value** | **P value<sup>a</sup>** |
| Missense | 1 | 2.4 | 0.417 | 0.909 |

<sup>a</sup> P-values in bold survived Bonferroni multiple test correction for the 13 tests.

### Functional annotation analysis

Following the significant enrichments above, we investigated if the genes contributing to each of the neurodevelopmental phenotypes influence the same or different biological functions by generating five subsets of the MEF2C gene-set based; (i) genes associated with SZ (n=41), (ii) genes associated with IQ (n=37), (iii) genes associated with EA (n=109), (iv) genes containing LoF DNMs in ASD cases (n=45) and (v) genes containing missense DNMs in ID cases (n=25). GO analysis identified different terms to be significantly enriched for these gene subsets. These data are summarized in Table 3 and detailed in S5-9 Tables. Terms related to development of the nervous systems are enriched in analyses of all five subsets of genes. Terms related to synaptic signalling are strongly enriched for SZ and EA. The most significantly enriched terms for IQ map to pre- and post-synapse function. The most significantly enriched terms for EA map to the synaptic membrane and synaptic signalling. For ASD and ID genes, other significantly enriched terms map to cell morphogenesis.

**Table 3:** Enriched GO terms for genes in the MEF2C gene-set contributing to different phenotypes

| Gene Subset | # Genes | Category<sup>a</sup> | Selected Enriched Terms of Shared Ancestry (GO:IDs) |
| --- | --- | --- | --- |
| MEF2C-SZ | 41 | BP | neurogenesis (0022008), regulation of nervous system development (0051960), neuron differentiation (0030182), neuron development (0048666), trans-synaptic signalling (0099537), synaptic signalling (0099536) |
| MEF2C-IQ | 37 | CC | postsynapse (0098794), neuron to neuron synapse (0098984), presynaptic cytoskeleton (0099569), |
|  |  | BP | neuron differentiation (0030182), neuron development (0048666), trans-synaptic signalling (0099537) |
| MEF2C-EA | 109 | CC | synaptic membrane (0097060), integral component of synaptic membrane (0099699), plasma membrane region (0098590), intrinsic component of synaptic membrane (0099240) |
|  |  | BP | anterograde trans-synaptic signalling (0098916), trans-synaptic signalling (0099537), synaptic signalling (0099536), cell-cell signalling (0007267), nervous system development (0007399) |
| MEF2C-ASD | 45 | BP | cell projection morphogenesis (0048858), cell morphogenesis (0000902), cell part morphogenesis (0032990), cellular component morphogenesis (0032989), neuron differentiation (0030182), neuron development (0048666) |
| MEF2C-ID | 25 | BP | forebrain neuron fate commitment (0021877), neuron differentiation (0030182), cell morphogenesis (0000902), cell morphogenesis involved in differentiation (0000904), neuron development (0048666) |

<sup>a</sup> BP = biological process; CC = cellular compartment

### Cell type enrichment analysis

By performing enrichment analysis on gene expression data from scRNA-seq of the mouse nervous system, we sought to detect individual cell types that are enriched for genes that are differentially expressed upon cKO of MEF2C. We used two scRNA-seq data-sets that included 265 cell types from the mouse nervous system (28) and 565 cell types from the mouse brain (29). After Bonferroni correcting for the 830 cell types tested, we identified 31 significantly enriched cell types in both data-sets (62 cell types in total with p<6.03×10^−5^). These results are summarized in Table 4 and full details are provided in S9-10 Tables.

**Table 4:**
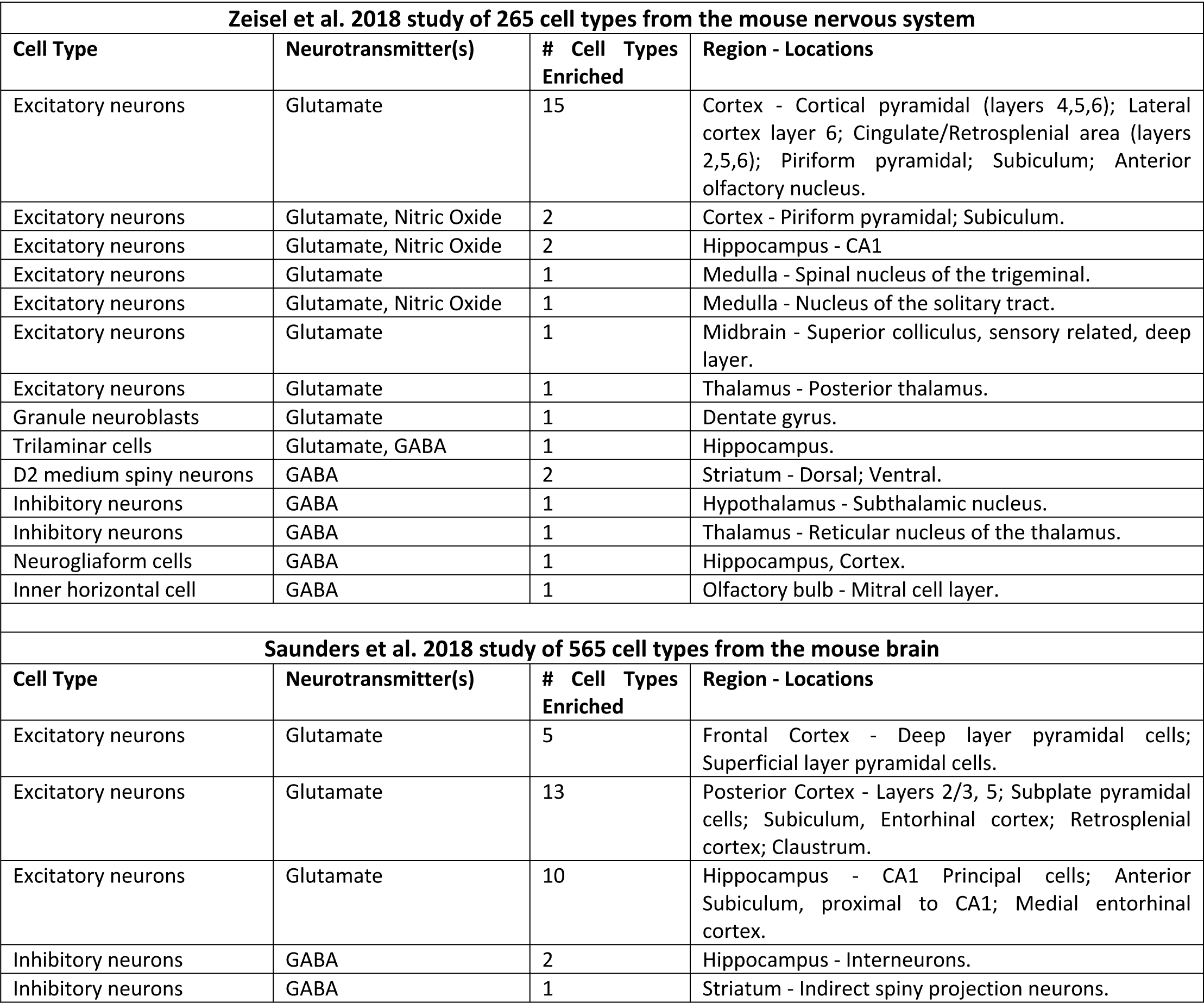
Enriched cell types from analysis of scRNA-seq data using the MEF2C gene-set

| **Zeisel et al. 2018 study of 265 cell types from the mouse nervous system** |  |  |  |
| --- | --- | --- | --- |
| **Cell Type** | **Neurotransmitter(s)** | **# Cell Types Enriched** | **Region - Locations** |
| Excitatory neurons | Glutamate | 15 | Cortex - Cortical pyramidal (layers 4,5,6); Lateral cortex layer 6; Cingulate/Retrosplenial area (layers 2,5,6); Piriform pyramidal; Subiculum; Anterior olfactory nucleus. |
| Excitatory neurons | Glutamate, Nitric Oxide | 2 | Cortex - Piriform pyramidal; Subiculum. |
| Excitatory neurons | Glutamate, Nitric Oxide | 2 | Hippocampus - CA1 |
| Excitatory neurons | Glutamate | 1 | Medulla - Spinal nucleus of the trigeminal. |
| Excitatory neurons | Glutamate, Nitric Oxide | 1 | Medulla - Nucleus of the solitary tract. |
| Excitatory neurons | Glutamate | 1 | Midbrain - Superior colliculus, sensory related, deep layer. |
| Excitatory neurons | Glutamate | 1 | Thalamus - Posterior thalamus. |
| Granule neuroblasts | Glutamate | 1 | Dentate gyrus. |
| Trilaminar cells | Glutamate, GABA | 1 | Hippocampus. |
| D2 medium spiny neurons | GABA | 2 | Striatum - Dorsal; Ventral. |
| Inhibitory neurons | GABA | 1 | Hypothalamus - Subthalamic nucleus. |
| Inhibitory neurons | GABA | 1 | Thalamus - Reticular nucleus of the thalamus. |
| Neurogliaform cells | GABA | 1 | Hippocampus, Cortex. |
| Inner horizontal cell | GABA | 1 | Olfactory bulb - Mitral cell layer. |
| **Saunders et al. 2018 study of 565 cell types from the mouse brain** |  |  |  |
| **Cell Type** | **Neurotransmitter(s)** | **# Cell Types Enriched** | **Region - Locations** |
| Excitatory neurons | Glutamate | 5 | Frontal Cortex - Deep layer pyramidal cells; Superficial layer pyramidal cells. |
| Excitatory neurons | Glutamate | 13 | Posterior Cortex - Layers 2/3, 5; Subplate pyramidal cells; Subiculum, Entorhinal cortex; Retrosplenial cortex; Claustrum. |
| Excitatory neurons | Glutamate | 10 | Hippocampus - CA1 Principal cells; Anterior Subiculum, proximal to CA1; Medial entorhinal cortex. |
| Inhibitory neurons | GABA | 2 | Hippocampus - Interneurons. |
| Inhibitory neurons | GABA | 1 | Striatum - Indirect spiny projection neurons. |

Results were consistent between the two scRNA-seq data-sets where only neurons were enriched and these were predominantly glutamatergic excitatory neurons in the cortex, e.g. deep layer pyramidal cells and cell in the retrosplenial cortex, entorhinal cortex and subiculum. These are the specific cell types in the adult mouse brain that are likely to be most impacted by cKO of Mef2c during embryonic development. It is possible that the enrichment detected here could be due to the MEF2C gene-set representing a set of genes that are expressed in the somatosensory cortex, which is their origin. To investigate this we extracted a similarly-sized gene-set from the Harrington et al. (13) study that included genes that were expressed in the somatosensory cortex but were not differentially expressed upon cKO of MEF2C. These analyses identified just 6 of 830 cell types that were significantly enriched after multiple test correction (data not shown), ~10% of the number of cell types identified from analyses of the set of DEGs, and indicating that these cell type enrichments for the MEF2C DEGs are not simply a function of analyses of cortically-expressed genes. Although these DEGs are from analysis of the cortex, they are also enriched in other cell types: excitatory neurons in the hippocampus, e.g. cells in the cornu ammonis 1 (CA1) subfield, and GABAergic inhibitory neurons in hippocampus and striatum, e.g. dopamine D2 receptor expressing medium spiny neurons (MSNs; also called indirect spiny projection neurons; Table 4). Overall, our results are consistent with previous cell type enrichment analysis of GWAS data that highlights the relevant cell types for both SZ and IQ to be cortical somatosensory pyramidal cells, hippocampal CA1 pyramidal cells and striatal MSNs (22, 30). Thus, the cell types enriched for SZ- and IQ-associated genes are those cell types that are enriched for genes that are differentially expressed as a result of cKO of *Mef2C* during neurodevelopment.

## Discussion

Across the MEF2C DEGs, there is a difference in terms of the type of genetic variant and the type of phenotype it associates with. The missense and LoF DNMs in ID and ASD cases are likely to disrupt or ablate one copy of a gene and are associated here with these severe early age-of-onset neurodevelopmental disorders. Common SNPs of weak effect underpin the enrichment of DEGs in genes associated with SZ, IQ and EA and these SNPs are likely to have subtle effects on the regulation and/or expression of these genes. Gene ontology analysis identifies both consistencies and differences between the biological processes that are enriched for DEGs contributing to the different phenotypes. Terms related to development of neurons are highly enriched for genes contributing to the five phenotypes. Terms related to synaptic function feature strongly in the analysis of SZ, IQ and EA but less so for ASD and not at all for ID. Terms related to cell morphogenesis feature strongly in the analysis of ASD and ID but do not rank as high for SZ, IQ and EA. Overall, it appears that the phenotypic effect of different genetic variants depends on (i) the effect of the variant on gene function, with larger effect mutations associating with early onset illness and (ii) the type of biological process impacted, with synaptic function being affected by genetic variants associating with SZ, IQ and EA. The latter finding above is consistent with previous analysis showing that the DEGs that are down-regulated upon cKO of MEF2C are involved in synaptic transmission (13) and our analysis showing that these down-regulated genes show greater enrichment for genes associated with SZ, IQ and EA than the up-regulated genes. A caveat to our analysis of GWAS data is that with a smaller sample size, the ASD GWAS may be under-powered to detect an enrichment of *MEF2C* DEGs.

Analysis of the *MEF2C* locus in GWAS indicates that although the signals for SZ and for IQ and EA co-localize to the 3’ end and downstream region of the gene, the SNPs associated with SZ risk and cognitive function are not in high LD with each other. For the IQ/EA association signal, one eQTL study in *post-mortem* adult cerebellum shows that the alleles associated with reduced IQ and EA are associated with increased expression of *MEF2C*. This eQTL was not consistently detected across multiple eQTL analysis of different brain regions and is contrary to data that shows upregulation of Mef2c in prefrontal projection neurons in mice resulted in enhanced cognitive performance (18). Ultimately, these are small effects and we do yet not know what effect these SZ- or IQ/EA-associated variants at *MEF2C* have on gene function in different brain regions and at different stages of development, and what the knock-on effects are on genes influenced by MEF2C.

scRNA-seq has delivered empirical taxonomies of brain cell types (29). Analysis of these data has allowed us to exploit the RNA-seq data to identify the individual cell types likely impacted by cKO of *Mef2c*. Genes that are differentially expressed upon cKO of *Mef2c* are specifically highly expressed across a large number of different cell types. As expected, based on the cell type targeted for cKO of *Mef2c* in the animal model, the majority of cell types affected are glutamatergic excitatory neurons. The advantage of these large scRNA-seq resources is that we can identify some of the specific regions and cell types affected. A number of the regional locations were identified in both analyses of scRNA-seq data and have previously been implicated in neurodevelopmental disorders. Transcriptomic analysis of dorsolateral prefrontal cortex layer 3 and layer 5 pyramidal cells have identified changes in gene expression between SZ cases and controls (31). LoF mutations in transcription factor T-brain-1 (TBR1) have been reported in ASD patients and conditional Tbr1 deletion during late mouse gestation in cortical layer 6 neurons affects processes such as dendritic patterning and synaptogenesis (32). The retrosplenial cortex, a sub-region of the cingulate cortex, exhibited impaired function using a semantic memory task (33) and showed reduced grey matter volumes (34) in SZ patients. The common set of brain regions that underlies the default mode network, autobiographical memory, prospection, navigation, and theory of mind may include the retrosplenial cortex (35). Resting state connectivity of the retrosplenial cortex is altered in SZ patients, with connectivity differences observed to brain areas associated with language processing, suggesting a role for the retrosplenial cortex in the aetiology of hallucinations (36). In ASD, resting state overconnectivity between the anterior insula and the retrosplenial cortex is associated with internalizing symptoms (anxiety, depression, social withdrawal) (37)).

Two limitations of our study are: (i) The cKO of *Mef2c* only occurred in differentiated forebrain excitatory neurons during embryonic development and RNA-seq was only performed on the somatosensory cortex, thus we are not getting a whole-brain picture of what gene expression, biological pathways and cell types may be influenced if MEF2C was manipulated in all tissues. If KO of *Mef2c* results in altered expression of a similar set of genes in hippocampus and striatum, our scRNA-seq analysis suggests that glutamatergic excitatory neurons and GABAergic inhibitory neurons in the hippocampus, and GABAergic inhibitory D2 medium spiny neurons in the striatum would be affected. Such results would be consistent with the balance of excitatory and inhibitory synapses in the developing brain being an element of neurodevelopmental disease. (ii) RNA-seq was only performed on the adult mouse, thus we are not getting a picture of the effect cKO of *Mef2c* in the developing brain. We acknowledge that a set of DEGs from embryonic tissue would give us a better insight into the molecular mechanisms of neurodevelopmental phenotypes and how they are influenced by *Mef2c*. But what we are getting is insight into the molecular consequences of embryonic cKO of *Mef2c* on the adult brain, and importantly this cKO of *Mef2c* during neurodevelopment did cause the behavioural phenotypes in adult mice related to memory, social behaviour, reward-related behaviours and repetitive motor behaviours that are consistent with neurodevelopmental disorders (13). What underpins these behavioural phenotypes in the mouse model is a reduction in cortical network activity, partly as a result of an increase in inhibitory and a decrease in excitatory synaptic transmission (13). We find that these MEF2C DEGs are enriched for genes associated with SZ, IQ and EA, based on analysis of common variants, and are enriched for genes that contribute to ASD and ID, based on analysis of rare mutations. Therefore, it is reasonable to conclude that the specific glutamatergic excitatory neurons in the cortex, which we find to be enriched for these MEF2C DEGs, are fundamental to the pathophysiology of neurodevelopmental disorders.

## Materials and Methods

### Ethics statement

Data were directly downloaded from published studies and no additional ethics approval was needed. Each study is referenced and details on ethics approval are available in each manuscript.

### MEF2C gene-set

There were 1076 DEGs reported in *Mef2c* cKO mice compared with controls with a log2 fold change >0.3 and at a FDR <0.05 (478 genes with increased expression level (up-regulated) in cKO mice and 598 genes with decreased expression level (down-regulated)) (13). This mouse DEG list was converted to human orthologue genes using Ensembl Biomart https://www.ensembl.org/biomart) or using with NCBI Gene database (https://www.ncbi.nlm.nih.gov/gene) if Biomart failed to find an orthologue. This resulted in a gene-set of 1,055 genes (465 up-regulated and 590 down-regulated; S1 Table). Sets of ‘brain-expressed’ genes (n=14,243) and ‘brain-elevated’ genes (n=1,424) were sourced from the Human Protein Atlas (https://www.proteinatlas.org/humanproteome/brain) and used as covariates in analyses. Brain-elevated genes are those that show an elevated expression in brain compared to other tissue types.

### GWAS data

This MEF2C gene-set was tested for enrichment of genes associated with different neurodevelopmental phenotypes using GWAS summary statistics for SZ (40,675 cases and 64,643 controls) (20), ASD (18,381 cases and 27,969 controls) (19), IQ (269,867 individuals) (22) and EA (1,131,881 individuals) (24). For control purposes, we tested for enrichment using GWAS summary statistics for ten other phenotypes. These were attention deficit hyperactivity disorder (ADHD (38)), bipolar disorder (BPD (39)), major depressive disorder (MDD (40)), obsessive-compulsive disorder (OCD (41)), Alzheimer’s disease (AD (42)), Crohn’s disease (CD (43)), height (44), stroke (45), type 2 diabetes (T2D (46)) and inflammatory bowel disease (IBD (47)).

### eQTL data

Data from the following were used to identify eQTLs at *MEF2C*: PsychENCODE, adult human cortex (48) (http://resource.psychencode.org/); Genotype-Tissue Expression (GTEx), multiple brain regions (https://gtexportal.org/); BRAINEAC, multiple brain regions (http://www.braineac.org); BrainSeq, hippocampus and dorsolateral prefrontal cortex (http://eqtl.brainseq.org/phase2/); O’Brien et al. (49), human fetal brain. HaploReg v4.1 (50) (https://pubs.broadinstitute.org/mammals/haploreg/haploreg.php), links to eQTLs from multiple sources.

### MAGMA gene-set analysis

A gene-set analysis (GSA) is a statistical method for simultaneously analysing multiple genetic markers in order to determine their joint effect. We performed GSA using MAGMA (51) and summary statistics from various GWAS. A MAGMA analysis involved three steps. First, in the annotation step we mapped SNPs with available GWAS results on to genes (GRCh37/hg19 start-stop coordinates +/-20kb). Second, in the gene analysis step we computed gene P values for each GWAS dataset. This gene analysis is based on a multiple linear principal components regression model that accounts for LD between SNPs. The European panel of the 1000 Genomes data was used as a reference panel for LD. Third, a competitive GSA based on the gene P values, also using a regression structure, was used to test if the genes in a gene-set were more strongly associated with either phenotype than other genes in the genome. The MHC region is strongly associated in the SZ GWAS data. This region contains high LD and the association signal has been attributed to just a small number of independent variants (52). However, MAGMA still identifies a very large number of associated genes despite factoring in the LD information. To avoid the excessive number of associated genes biasing the MAGMA GSA, we excluded all genes within the MHC region from our GSA of SZ. MAGMA was chosen because it corrects for LD, gene size and gene density (potential confounders) and has significantly more power than other GSA tools (53). The second step of MAGMA analysis generates a P value for each gene. By Bonferroni correcting for the number of genes tested in each GWAS, we used these data to identify individual genes that genome-wide associated with SZ (n=499), IQ (n=647), and EA (n=1,278).

### Enrichment analysis for genes containing *de novo* mutations

Lists of genes harbouring DNMs identified in patients with SZ (n=1,024), ASD (n=3,985), ID (n=192) and in unaffected siblings (n=1,995) and controls (n=54) based on exome sequencing of trios were sourced from Genovese et al. (54). DNMs were categorized as silent, missense and loss-of-function (LoF). We tested for enrichment of our MEF2C gene-set in these gene lists using denovolyzeR (55). DenovolyzeR is an R package for the statistical analysis of DNMs. For each gene, denovolyzeR derives the expected number of DNMs in a given population based on the mutability of the gene and the number of trios sequenced. It then compares the observed number of DNMs against expectation using a Poisson framework to determine whether there is an excess of DNMs in a given gene or gene-set (55).

### Enrichment analysis of single cell transcriptomic data from the mouse nervous system

The expression weighted cell-type enrichment (EWCE) R package (https://github.com/NathanSkene/EWCE) represents a method to statistically evaluate if a set of genes (e.g. our MEF2C gene-set) has higher expression within a particular cell type than can be reasonably expected by chance (56). The probability distribution for this is estimated by randomly generating gene-sets of equal length from a set of background genes. We used scRNA-seq data from 19 regions across the central and peripheral nervous system of post-natal day (P) 12-30 and 6-8 week old mice (28) and from nine regions of the adult (P60-70) mouse brain (29). Tissue analysed by Zeisel et al. (28) included the anterior, middle and posterior cortex, hippocampus, amygdala, striatum, thalamus, hypothalamus, midbrain, cerebellum, dorsal root ganglion, myenteric plexus, submucosal plexus, muscle layer, sympathetic chain, spinal cord, pons, medulla and olfactory bulb. This resulted in 265 cell clusters represented by 160,796 sc transcriptomes. These expression datasets were sourced from http://mousebrain.org/. Regions analysed by Saunders et al. (29) included the frontal and posterior cortex, hippocampus, thalamus, cerebellum, striatum, globus pallidus externus and nucleus basalis, entopeduncular nucleus and subthalamic nucleus, and substantia nigra and ventral tagmental area. This resulted in 565 transcriptionally distinct populations from 690,000 individual cells. These expression datasets were sourced from http://dropviz.org/. We used the EWCE package to test whether MEF2C DEGs are enriched in any particular cell type from the scRNA-seq data. The analysis is carried out in three steps. First, the relevant scRNA-seq data is loaded. Next, a target gene-set and suitable background set is chosen. Finally, a bootstrapping enrichment test is run, controlling for transcript length and GC-content. The package controls for these by selecting bootstrap lists containing genes with similar properties to genes in the target list. For our analysis we ran 100,000 permutations. The package can convert gene IDs and provides a background set of mouse/human genes from Biomart. Genes in the target or background set without mouse/human orthologs were dropped.

### Functional annotation

ConsensusPathDB-human (http://cpdb.molgen.mpg.de/) was used to perform overrepresentation analysis of gene-sets and we report on enriched Gene Ontology (GO) terms (57).

## Supplementary Figure Legends

S1 Figure A-D: GWAS results at MEF2C in ASD (A), EA (B), IQ (C) and SZ (D). Chromosomal position (Mb) is plotted on the x-axis and -log10 GWAS p-values for SNPs are plotted on left y-axis. The blue line denotes regional recombination rates (cMMb) plotted on the right y-axis. The index SNP, which is the SNP most associated with the phenotype, is represented by a purple diamond. SNPs (dots) are colour coded based on degree of LD to the single index SNP as represented by r2. Legend for r2 is shown in upper right corner. LocusZoom (http://locuszoom.org/) was used for visualization of GWAS results at the MEF2C locus for the different phenotypes.

## Supplementary Table Information

S1 Table: MEF2C gene-set containing DEGs from somatosensory cortex following cKO of Mef2c in mice

S2 Table: Conditional MAGMA GSA for SZ, IQ and EA where results survived correction in Table 1

S3 Table: MAGMA GSA of other phenotypes

S4 Table: GO analysis of the subset of MEF2C DEGs that are associated with SZ

S5 Table: GO analysis of the subset of MEF2C DEGs that are associated with IQ

S6 Table: GO analysis of the subset of MEF2C DEGs that are associated with EA

S7 Table: GO analysis of the subset of MEF2C DEGs that contain LoF variants reported in ASD

S8 Table: GO analysis of the subset of MEF2C DEGs that contain missense variants reported in ID

S9 Table: EWCE analysis of MEF2C DEGs in the Zeisel et al. 2018 scRNA-seq data for 265 cell types

S10 Table: EWCE analysis of MEF2C DEGs in the Saunders et al. 2018 scRNA-seq data for 565 cell types

## References

1. Li H, Radford JC, Ragusa MJ, Shea KL, McKercher SR, Zaremba JD, et al. Transcription factor MEF2C influences neural stem/progenitor cell differentiation and maturation in vivo. Proceedings of the National Academy of Sciences of the United States of America. 2008;105(27):9397–402.

2. Potthoff MJ, Olson EN. MEF2: a central regulator of diverse developmental programs. Development (Cambridge, England). 2007;134(23):4131–40.

3. Consortium U. UniProt: a hub for protein information. Nucleic acids research. 2015;43(Database issue):D204–12.

4. Lyons GE, Micales BK, Schwarz J, Martin JF, Olson EN. Expression of mef2 genes in the mouse central nervous system suggests a role in neuronal maturation. The Journal of neuroscience: the official journal of the Society for Neuroscience. 1995;15(8):5727–38.

5. Haberland M, Arnold MA, McAnally J, Phan D, Kim Y, Olson EN. Regulation of HDAC9 gene expression by MEF2 establishes a negative-feedback loop in the transcriptional circuitry of muscle differentiation. Molecular and cellular biology. 2007;27(2):518–25.

6. Lin Q, Schwarz J, Bucana C, Olson EN. Control of mouse cardiac morphogenesis and myogenesis by transcription factor MEF2C. Science (New York, NY). 1997;276(5317):1404–7.

7. Adachi M, Lin PY, Pranav H, Monteggia LM. Postnatal Loss of Mef2c Results in Dissociation of Effects on Synapse Number and Learning and Memory. Biological psychiatry. 2016;80(2):140–8.

8. Chen YC, Kuo HY, Bornschein U, Takahashi H, Chen SY, Lu KM, et al. Foxp2 controls synaptic wiring of corticostriatal circuits and vocal communication by opposing Mef2c. Nature neuroscience. 2016;19(11):1513–22.

9. Tu S, Akhtar MW, Escorihuela RM, Amador-Arjona A, Swarup V, Parker J, et al. NitroSynapsin therapy for a mouse MEF2C haploinsufficiency model of human autism. Nature communications. 2017;8(1):1488.

10. Barbosa AC, Kim MS, Ertunc M, Adachi M, Nelson ED, McAnally J, et al. MEF2C, a transcription factor that facilitates learning and memory by negative regulation of synapse numbers and function. Proceedings of the National Academy of Sciences of the United States of America. 2008;105(27):9391–6.

11. Lipton SA, Li H, Zaremba JD, McKercher SR, Cui J, Kang YJ, et al. Autistic phenotype from MEF2C knockout cells. Science (New York, NY). 2009;323(5911):208.

12. Cho EG, Zaremba JD, McKercher SR, Talantova M, Tu S, Masliah E, et al. MEF2C enhances dopaminergic neuron differentiation of human embryonic stem cells in a parkinsonian rat model. PloS one. 2011;6(8):e24027.

13. Harrington AJ, Raissi A, Rajkovich K, Berto S, Kumar J, Molinaro G, et al. MEF2C regulates cortical inhibitory and excitatory synapses and behaviors relevant to neurodevelopmental disorders. Elife. 2016;5.

14. Paciorkowski AR, Traylor RN, Rosenfeld JA, Hoover JM, Harris CJ, Winter S, et al. MEF2C Haploinsufficiency features consistent hyperkinesis, variable epilepsy, and has a role in dorsal and ventral neuronal developmental pathways. Neurogenetics. 2013;14(2):99–111.

15. Rocha H, Sampaio M, Rocha R, Fernandes S, Leao M. MEF2C haploinsufficiency syndrome: Report of a new MEF2C mutation and review. Eur J Med Genet. 2016;59(9):478–82.

16. Mikhail FM, Lose EJ, Robin NH, Descartes MD, Rutledge KD, Rutledge SL, et al. Clinically relevant single gene or intragenic deletions encompassing critical neurodevelopmental genes in patients with developmental delay, mental retardation, and/or autism spectrum disorders. American journal of medical genetics Part A. 2011;155A(10):2386–96.

17. Le Meur N, Holder-Espinasse M, Jaillard S, Goldenberg A, Joriot S, Amati-Bonneau P, et al. MEF2C haploinsufficiency caused by either microdeletion of the 5q14.3 region or mutation is responsible for severe mental retardation with stereotypic movements, epilepsy and/or cerebral malformations. Journal of medical genetics. 2010;47(1):22–9.

18. Mitchell AC, Javidfar B, Pothula V, Ibi D, Shen EY, Peter CJ, et al. MEF2C transcription factor is associated with the genetic and epigenetic risk architecture of schizophrenia and improves cognition in mice. Mol Psychiatry. 2017.

19. Grove J, Ripke S, Als TD, Mattheisen M, Walters RK, Won H, et al. Identification of common genetic risk variants for autism spectrum disorder. Nature genetics. 2019;51(3):431–44.

20. Pardinas AF, Holmans P, Pocklington AJ, Escott-Price V, Ripke S, Carrera N, et al. Common schizophrenia alleles are enriched in mutation-intolerant genes and in regions under strong background selection. Nature genetics. 2018;50(3):381–9.

21. Sniekers S, Stringer S, Watanabe K, Jansen PR, Coleman JRI, Krapohl E, et al. Genome-wide association meta-analysis of 78,308 individuals identifies new loci and genes influencing human intelligence. Nature genetics. 2017;49(7):1107–12.

22. Savage JE, Jansen PR, Stringer S, Watanabe K, Bryois J, de Leeuw CA, et al. Genome-wide association meta-analysis in 269,867 individuals identifies new genetic and functional links to intelligence. Nature genetics. 2018;50(7):912–9.

23. Davies G, Lam M, Harris SE, Trampush JW, Luciano M, Hill WD, et al. Study of 300,486 individuals identifies 148 independent genetic loci influencing general cognitive function. Nature communications. 2018;9(1):2098.

24. Lee JJ, Wedow R, Okbay A, Kong E, Maghzian O, Zacher M, et al. Gene discovery and polygenic prediction from a genome-wide association study of educational attainment in 1.1 million individuals. Nature genetics. 2018;50(8):1112–21.

25. Okbay A, Beauchamp JP, Fontana MA, Lee JJ, Pers TH, Rietveld CA, et al. Genome-wide association study identifies 74 loci associated with educational attainment. Nature. 2016;533(7604):539–42.

26. Zou F, Chai HS, Younkin CS, Allen M, Crook J, Pankratz VS, et al. Brain expression genome-wide association study (eGWAS) identifies human disease-associated variants. PLoS Genet. 2012;8(6):e1002707.

27. Gandal MJ, Zhang P, Hadjimichael E, Walker RL, Chen C, Liu S, et al. Transcriptome-wide isoform-level dysregulation in ASD, schizophrenia, and bipolar disorder. Science. 2018;362(6420):eaat8127.

28. Zeisel A, Hochgerner H, Lönnerberg P, Johnsson A, Memic F, van der Zwan J, et al. Molecular Architecture of the Mouse Nervous System. Cell. 2018;174(4):999–1014.e22.

29. Saunders A, Macosko EZ, Wysoker A, Goldman M, Krienen FM, de Rivera H, et al. Molecular Diversity and Specializations among the Cells of the Adult Mouse Brain. Cell. 2018;174(4):1015–30 e16.

30. Skene NG, Bryois J, Bakken TE, Breen G, Crowley JJ, Gaspar HA, et al. Genetic identification of brain cell types underlying schizophrenia. Nature Genetics. 2018;50(6):825–33.

31. Arion D, Huo Z, Enwright JF, Corradi JP, Tseng G, Lewis DA. Transcriptome Alterations in Prefrontal Pyramidal Cells Distinguish Schizophrenia From Bipolar and Major Depressive Disorders. Biological psychiatry. 2017;82(8):594–600.

32. Fazel Darbandi S, Robinson Schwartz SE, Qi Q, Catta-Preta R, Pai EL, Mandell JD, et al. Neonatal Tbr1 Dosage Controls Cortical Layer 6 Connectivity. Neuron. 2018;100(4):831–45 e7.

33. Tendolkar I, Weis S, Guddat O, Fernandez G, Brockhaus-Dumke A, Specht K, et al. Evidence for a dysfunctional retrosplenial cortex in patients with schizophrenia: a functional magnetic resonance imaging study with a semantic-perceptual contrast. Neuroscience letters. 2004;369(1):4–8.

34. Mitelman SA, Shihabuddin L, Brickman AM, Hazlett EA, Buchsbaum MS. Volume of the cingulate and outcome in schizophrenia. Schizophr Res. 2005;72(2-3):91–108.

35. Spreng RN, Mar RA, Kim AS. The common neural basis of autobiographical memory, prospection, navigation, theory of mind, and the default mode: a quantitative meta-analysis. Journal of cognitive neuroscience. 2009;21(3):489–510.

36. Bluhm RL, Miller J, Lanius RA, Osuch EA, Boksman K, Neufeld RW, et al. Retrosplenial cortex connectivity in schizophrenia. Psychiatry Res. 2009;174(1):17–23.

37. Hogeveen J, Krug MK, Elliott MV, Solomon M. Insula-Retrosplenial Cortex Overconnectivity Increases Internalizing via Reduced Insight in Autism. Biological psychiatry. 2018;84(4):287–94.

38. Demontis D, Walters RK, Martin J, Mattheisen M, Als TD, Agerbo E, et al. Discovery of the first genome-wide significant risk loci for attention deficit/hyperactivity disorder. Nature Genetics. 2019;51(1):63–75.

39. Stahl EA, Breen G, Forstner AJ, McQuillin A, Ripke S, Trubetskoy V, et al. Genome-wide association study identifies 30 loci associated with bipolar disorder. Nature genetics. 2019;51(5):793–803.

40. Wray NR, Ripke S, Mattheisen M, Trzaskowski M, Byrne EM, Abdellaoui A, et al. Genome-wide association analyses identify 44 risk variants and refine the genetic architecture of major depression. Nature genetics. 2018;50(5):668–81.

41. International Obsessive Compulsive Disorder Foundation Genetics C, Studies OCDCGA, Arnold PD, Askland KD, Barlassina C, Bellodi L, et al. Revealing the complex genetic architecture of obsessive–compulsive disorder using meta-analysis. Molecular Psychiatry. 2017;23:1181.

42. Lambert JC, Ibrahim-Verbaas CA, Harold D, Naj AC, Sims R, Bellenguez C, et al. Meta-analysis of 74,046 individuals identifies 11 new susceptibility loci for Alzheimer’s disease. Nature genetics. 2013;45(12):1452–8.

43. Franke A, McGovern DP, Barrett JC, Wang K, Radford-Smith GL, Ahmad T, et al. Genome-wide meta-analysis increases to 71 the number of confirmed Crohn’s disease susceptibility loci. Nature genetics. 2010;42(12):1118–25.

44. Wood AR, Esko T, Yang J, Vedantam S, Pers TH, Gustafsson S, et al. Defining the role of common variation in the genomic and biological architecture of adult human height. Nature genetics. 2014;46:1173.

45. Traylor M, Farrall M, Holliday EG, Sudlow C, Hopewell JC, Cheng YC, et al. Genetic risk factors for ischaemic stroke and its subtypes (the METASTROKE collaboration): a meta-analysis of genome-wide association studies. The Lancet Neurology. 2012;11(11):951–62.

46. Mahajan A, Go MJ, Zhang W, Below JE, Gaulton KJ, Ferreira T, et al. Genome-wide trans-ancestry meta-analysis provides insight into the genetic architecture of type 2 diabetes susceptibility. Nature genetics. 2014;46(3):234–44.

47. Liu JZ, van Sommeren S, Huang H, Ng SC, Alberts R, Takahashi A, et al. Association analyses identify 38 susceptibility loci for inflammatory bowel disease and highlight shared genetic risk across populations. Nature genetics. 2015;47(9):979–86.

48. Wang D, Liu S, Warrell J, Won H, Shi X, Navarro FCP, et al. Comprehensive functional genomic resource and integrative model for the human brain. Science. 2018;362(6420):eaat8464.

49. O’Brien HE, Hannon E, Hill MJ, Toste CC, Robertson MJ, Morgan JE, et al. Expression quantitative trait loci in the developing human brain and their enrichment in neuropsychiatric disorders. Genome Biology. 2018;19(1):194.

50. Ward LD, Kellis M. HaploReg: a resource for exploring chromatin states, conservation, and regulatory motif alterations within sets of genetically linked variants. Nucleic Acids Res. 2012;40(Database issue):D930–4.

51. de Leeuw CA, Mooij JM, Heskes T, Posthuma D. MAGMA: Generalized Gene-Set Analysis of GWAS Data. PLOS Computational Biology. 2015;11(4):e1004219.

52. Sekar A, Bialas AR, de Rivera H, Davis A, Hammond TR, Kamitaki N, et al. Schizophrenia risk from complex variation of complement component 4. Nature. 2016;530(7589):177–83.

53. de Leeuw CA, Neale BM, Heskes T, Posthuma D. The statistical properties of gene-set analysis. Nat Rev Genet. 2016;17(6):353–64.

54. Genovese G, Fromer M, Stahl EA, Ruderfer DM, Chambert K, Landen M, et al. Increased burden of ultra-rare protein-altering variants among 4,877 individuals with schizophrenia. Nature neuroscience. 2016;19(11):1433–41.

55. Ware JS, Samocha KE, Homsy J, Daly MJ. Interpreting de novo Variation in Human Disease Using denovolyzeR. Current protocols in human genetics. 2015;87:7.25.1–7..15.

56. Skene NG, Grant SG. Identification of Vulnerable Cell Types in Major Brain Disorders Using Single Cell Transcriptomes and Expression Weighted Cell Type Enrichment. Front Neurosci. 2016;10:16.

57. Herwig R, Hardt C, Lienhard M, Kamburov A. Analyzing and interpreting genome data at the network level with ConsensusPathDB. Nat Protocols. 2016;11(10):1889–907.

